# Higher-order correction of persistent batch effects in correlation networks

**DOI:** 10.1101/2023.12.28.573533

**Authors:** Soel Micheletti, Daniel Schlauch, John Quackenbush, Marouen Ben Guebila

**Author notes:** Co-first authors (Soel Micheletti, Daniel Schlauch). Co-senior authors (John Quackenbush, Marouen Ben Guebila).

## Abstract

Systems biology methods often rely on correlations in gene expression profiles to infer co-expression networks, commonly used as input for gene regulatory network inference or to identify functional modules of co-expressed or co-regulated genes. While systematic biases, including batch effects, are known to induce spurious associations and confound differential gene expression analyses (DE), the impact of batch effects on gene co-expression has not been fully explored. Methods have been developed to adjust expression values, ensuring conditional independence of mean and variance from batch or other covariates for each gene. These adjustments have been shown to improve the fidelity of DE analysis. However, these methods do not address the potential for spurious differential co-expression (DC) between groups. Consequently, uncorrected, artifactual DC can skew the correlation structure, leading network inference methods that use gene co-expression to identify false, nonbiological associations, even when the input data is corrected using standard batch correction.

In this work, we demonstrate the persistence of confounders in covariance after standard batch correction using synthetic and real-world gene expression data examples. Subsequently, we introduce Co-expression Batch Reduction Adjustment (COBRA), a method for computing a batch-corrected gene co-expression matrix based on estimating a conditional covariance matrix. COBRA estimates a reduced set of parameters expressing the co-expression matrix as a function of the sample covariates, allowing control for continuous and categorical covariates. COBRA is computationally efficient, leveraging the inherently modular structure of genomic data to estimate accurate gene regulatory associations and facilitate functional analysis for high-dimensional genomic data.

## 1 Introduction

Batch effects in gene expression analyses arise because samples are typically collected and processed in different groups or batches due to logistical and practical reasons [Scherer, 2009, Lazar et al., 2013]. If not properly corrected, batch effects can introduce artifacts that obscure the true biology driving phenotypic differences [Leek et al., 2010]. Various methods exist to correct for batch effects in differential expression analysis, including ComBat [Johnson et al., 2007, Zhang et al., 2020], dChip [Li and Wong, 2003], as well as more general procedures like LIMMA [Ritchie et al., 2015] and SVA [Leek et al., 2012]. Although they differ in how they handle batch effects, each of these methods amount to a linear correction of batch effects including “location-scale correction” that adjusts the mean and variance of gene expression, addressing biases that can occur in differential expression analysis for both microarrays and RNA-Seq data [Conesa et al., 2016].

However, the factors distinguishing phenotypes go beyond the expression of particular genes, and involve genes that are coordinately activated or regulated in specific biological states. Network science methods have proven useful in modeling such complex relationships [Califano and Alvarez, 2017, Sinha et al., 2020], for instance by identifying differences between health and disease that cannot be uncovered using differential gene expression [Schlauch et al., 2017, Lopes-Ramos et al., 2021, Weighill et al., 2021]. Many of these methods consider the pairwise joint distribution of genes by creating and comparing co-expression networks [Hsu et al., 2015, Tesson et al., 2010, Langfelder and Horvath, 2008, Langfelder and Horvath, 2012, Southworth et al., 2009, Choi et al., 2005, Siska and Kechris, 2017, Yu et al., 2011, Amar et al., 2013], and they are able to identify functional groups of genes that are coordinately expressed in different biological states [Fuller et al., 2007, Lu and Keleş, 2023, Morabito et al., 2023]. These algorithms generally compute co-expression matrices following standard batch correction on gene expression data [Furlotte et al., 2011], and then compare the resulting networks between conditions. However, this is not sufficient to account for the possibility of batch-induced changes in correlation between genes, as existing batch correction methods act solely on the marginal distribution of each gene.

As an example, consider a system of two simulated genes, each of which is profiled across multiple samples organized in two batches (Figure 1-A). First, we consider a situation in which the expression of these genes is uncorrelated in both Batch A and Batch B, although with different batch-specific means and variances (upper left). We observe that batch correction using ComBat removes these differences and recovers their overall uncorrelated structure (upper right), so that the batches become, as expected, indistinguishable. Now, consider a situation where an artifactual correlation between the expression of gene 1 and gene 2 appears in Batch A but not Batch B (lower left); here ComBat batch correction centers the data from both batches, but fails to address the correlation structure that appeared in Batch 1 (lower right). This failure to address a higher order artifact can lead to spurious results in downstream analyses, such as inferring correlation-based networks.

**Figure 1:**
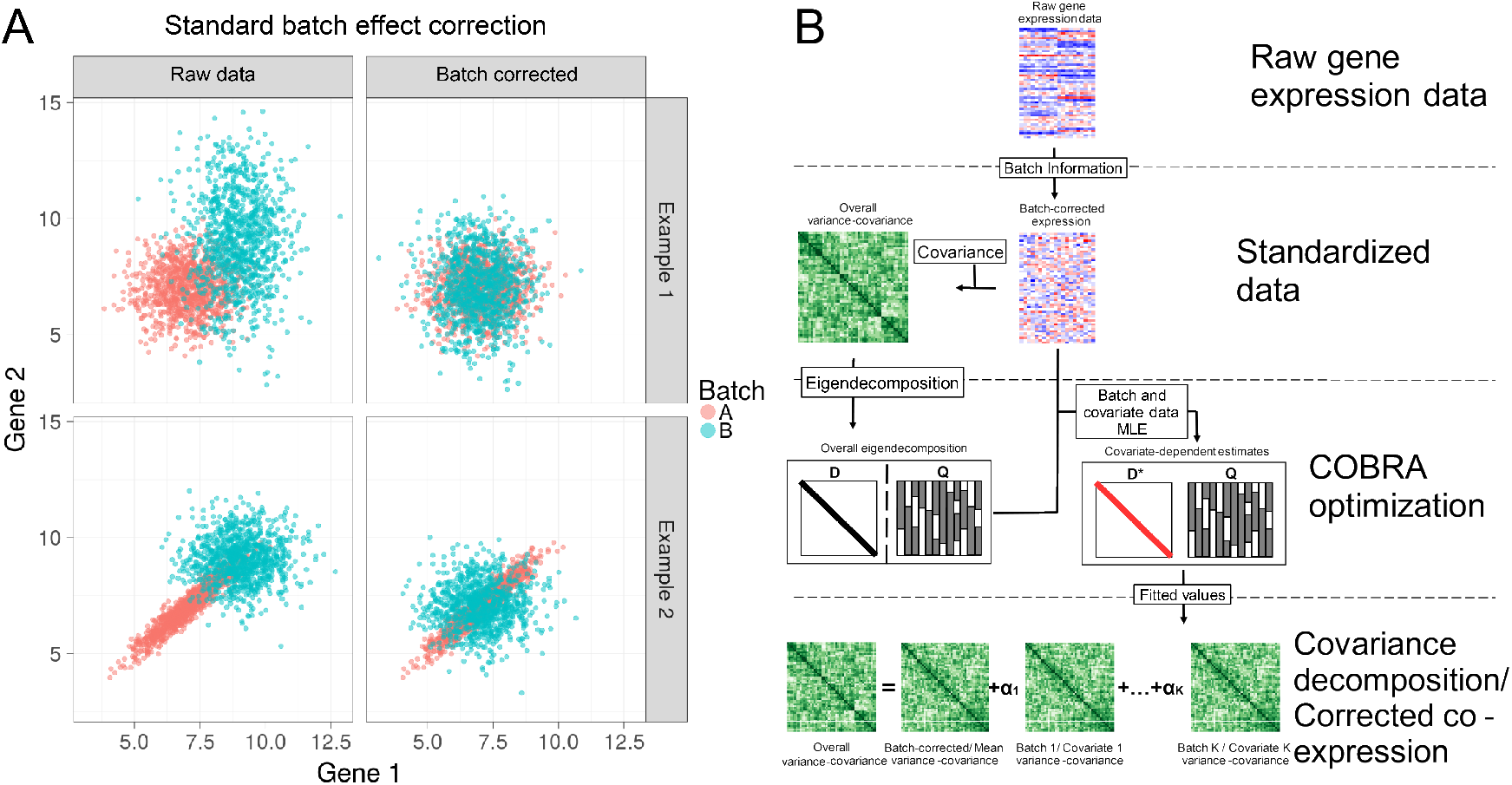
Identifying and correcting batch effects in correlation networks using COBRA. **A. Example demonstration of the impact of standard batch correction on co-expression.** An example of two simulated genes measured across many samples grouped into two batches. The top row shows a comparison of two conditionally independent genes, demonstrating that location-scale batch correction using ComBat, a standard batch correction method, appropriately removes the marginal dependence between them. The bottom row shows a similar set of simulated data. However, in this case, the two genes are artificially correlated in Batch A but not Batch B. The batch-dependent pattern of correlation persists after standard normalization. When comparing co-expression matrices, uncorrected batch-induced co-expression can bias co-expression analysis, making it challenging to tease out biological signal from technical effects. **B. The COBRA Workflow** begins with (1) normalizing the input gene expression data set, after which (2) the global co-expression matrix is calculated. (3) An eigendecomposition of the co-expression matrix is then computed. While retaining eigenvectors from this decomposition as a basis, we estimate “pseudo-eigenvalues” for each covariate, minimizing the error of the matrix reconstructed using these pseudo-eigenvalues compared to the original co-expression matrix. Finally, (4) the fitted pseudo-eigenvalues obtained from this estimation, in combination with the eigenvector matrix **Q**, are used for applications such as the inference of batch-corrected networks or covariate-specific co-expression analysis.

While some methods have addressed modeling the covariance matrix for small numbers of variables [Hoff and Niu, 2012, Zou et al., 2017], extending these approaches to handle higher-dimensional biological data, such as genome-scale transcriptional data, is nontrivial. First, the sample covariance matrix, 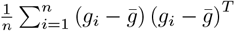, where *g*_*i*_ is a vector quantifying gene expression for sample *i* and *n* is the sample size, can be singular and therefore non positive-definite. Second, estimating the co-expression matrix involves inferring a large number of parameters. Given *p* genes, there are 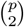 pairwise correlations, each of which must be a function of the number of covariates. In high-throughput gene expression studies, the number of genes *p* is typically much larger that the number samples *n*, and one often finds that co-expression in the same system does not consistently replicate across studies, likely because of the large numbers of parameters that must be estimated [Schlauch et al., 2017]. Imposing sparsity on the gene covariance matrix [Bien et al., 2011] or introducing a sparse precision matrix [Friedman et al., 2008] were suggested to address this issue, but this is not well suited for analyzing gene expression data where there is no *a priori* biologically motivated justification for excluding some subset of genes and even if there were, doing so can be computationally burdensome.

Co-expression Batch Reduction Adjustment (COBRA) addresses the problem of handling batch effects in estimating the gene expression covariance matrix by leveraging the modular nature of gene expression data. This approach effectively reduces the parameter space, estimating only 𝒪(*n*) variables for each covariate. To do this, COBRA uses information collected from many similarly expressed genes to perform a dimensionality reduction that allows us to estimate the gene co-expression matrix as a function of sample covariates. COBRA decomposes the co-expression matrix as a linear combination of components, one for each covariate, and uses a regression framework that allows continuous and categorical covariates to be included in an adjustment model (Figure 1-B). COBRA can be applied for batch correction, covariate-specific co-expression analysis, and to study the effect of various parameters on the observed patterns of co-expression.

## 2 Methods

### 2.1 Overview of the statistical approach

COBRA takes as input a gene expression matrix for *p* genes in *n* samples, and a design matrix *X* of dimension *n* × *q* containing a vector of *q* covariates for each sample. COBRA decomposes the co-expression matrix as a linear combination of components, one for each covariate in *X*, which can be used to correct for higher-moment batch effects in correlation matrices beyond biases in the mean and variance. By using the corrected co-expression matrix in network inference methods, we can effectively remove batch effects or identify covariate-specific co-expression patterns. In this section, after presenting our model and deriving an optimal estimator, we show how to correctly design *X* to tackle various problems.

### 2.2 Addressing persistent high-order batch effects using COBRA

Consider a set of *n* samples, for which we measure *q* covariates and the expression of *p* genes. Let 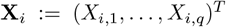 represent the covariates for sample *i*, and let **g**_*i*_ := (*g*_*i*,1_, …, *g*_*i,p*_)^*T*^ denote the gene expression values for sample *i* across the *p* genes. Moreover, let 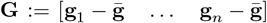 denote the zero-centered gene expression matrix, where 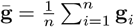 is the mean gene expression. To decompose the co-expression matrix, we model it as a function of the largest components of variation. This is coherent with the assumption that the underlying biology can be explained by a much smaller set of variance components among the 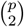 pairwise gene-gene relationships. This assumption allows us to preserve explainability by keeping all genes in the analysis, while also reflecting our understanding that it is groups of genes that act together to alter biological functions. Our approach to identify these components is to find a set of eigenvectors of the sample co-expression matrix **C** := **GG**^*T*^ which best explains the gene co-variation, using the fact that in high-dimensional settings, where *p > n*, the rank of *C* will be *r ≤ n*, resulting in at most *n* nonzero eigenvalues. Therefore, we only keep the eigenvectors of *C* corresponding to nonzero eigenvalues, substantially reducing the parameter space from 𝒪(*p*^2^) to 𝒪(*n*).

Specifically, we consider the reduced eigendecomposition **C** = **QDQ**^*T*^, where **D** is a diagonal matrix for the nonzero eigenvalues of **C**, and the columns of **Q** correspond to their respective eigenvectors. We then incorporate sample specific covariates in the design matrix to infer a diagonal matrix of “pseudo-eigenvalues”, quantifying the impact of each covariate on each nonzero eigenvalue. This inference is achieved by solving the following optimization problem:

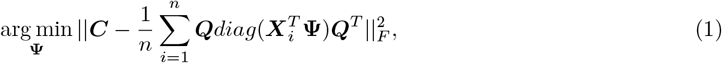

where **Ψ** is a *q* × *r* parameter matrix of coefficients that adjusts the eigenvalues as a function of the covariates to minimize the co-expression reconstruction error. For example, in the case of a single batch and in the absence of other experimental conditions, i.e. **X**_*i*_ = 1 for all *i* ∈ [*n*], then **Ψ** becomes identical to the vector of eigenvalues of the original covariance matrix. We show (Supplementary text 1) that this optimization problem in Equation 1 is equivalent to solving *r* linear regression problems admitting the following closed-form solution:

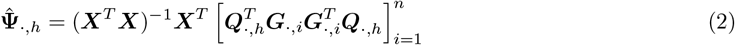

for all *h* ∈ [*r*]. Moreover, by including an intercept in the first column of the design matrix **X**, we achieve a global minimum in Equation 1 and derive the following the decomposition:

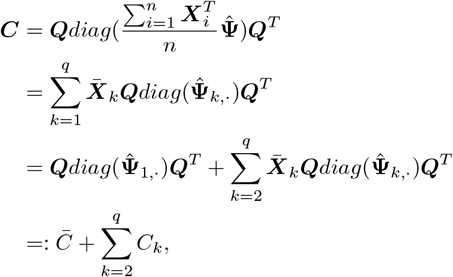

where 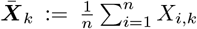. In other words, COBRA decomposes the co-expression matrix into a linear combination of components, one for each covariate in the design matrix. Depending on the choice of covariates in **X**, this can be used in a variety of different applications (Table 1) including batch correction, covariate-specific gene co-expression analysis, and understanding the association of particular variables with gene co-expression.

**Table 1:** COBRA use cases in correlation analyses.

| Goal | Design matrix | Output |
| --- | --- | --- |
| Batch correction | Column 1: intercept<br>Columns 2...$K$: Batch and covariates | $\bar{C}$ |
| Case/control comparison | Column 1: intercept<br>Column 2: 0 for control, 1 for case<br>Columns 3...$K$: Batch and covariates | $C_2$ |
| Covariate-specific co-expression | Column 1: intercept<br>Columns 2...$K$: covariates of interest | $C_k$ for $k \in [K]$ |

## 3 Results

To demonstrate batch effect control using COBRA, we conducted a benchmark in various applications such as case/control comparisons (Sections 3.1 and 3.2), batch effect removal (Sections 3.3 and 3.4), and extraction of covariate-specific co-expression (Sections 3.3 and 3.4). For each use case we provide examples on the type of covariates that can be included.

### 3.1 Improved co-expression estimates *in silico*

We conducted a simulation study in the presence of higher-order batch effects to investigate the performance of COBRA in identifying covariate-specific co-expression. We simulated gene expression for 4000 genes in each of 400 samples, where each sample was assigned to a batch group (A or B) and a treatment group (case or control). Based on the specific treatment-batch combination, we drew the gene expression levels for sample *i* from a multivariate Gaussian distribution with covariance Σ_*i*_ defined as:

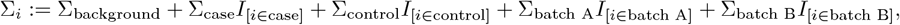

where the individuals components Σ_*j*_ for *j* ∈ {background, case, control, batch A, batch B} were designed to be sparse. This allowed us to reconstruct the ground truth: the nonzero pairs in Σ_case_ −Σ_control_ corresponded to real treatment effects, while the nonzero pairs in Σ_batch A_ − Σ_batch B_ corresponded to batch effects. To generate these covariance components, we created 10 modules with 400 genes each. The genes within each module exhibited a large absolute pairwise correlation sampled uniformly from the interval [−1, −0.5]∪[0.5, 1], and they were independent from genes outside their module. This resulted in a correlation matrix *C*_*k*_ for each module *M*_*k*_ (*k* ∈ [10]), where only 1% of the entries were nonzero. We assigned each module to background (*M*_1_, …, *M*_4_), case (*M*_5_ and *M*_6_), control (*M*_7_ and *M*_8_), batch A (*M*_9_), or batch B (*M*_10_), resulting in

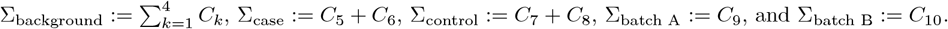

This simulation corresponds to a design matrix that has three variables representing three covariates: intercept for the average co-expression, batch (0 for Batch A, 1 for Batch B), and treatment (1 for case, 0 for control). We benchmarked the ability of COBRA to discriminate treatment effects, i.e. the nonzero pairs in *C*_case_ − *C*_control_, from the other pairs. As baselines we included the following batch correction methods: ComBat [Johnson et al., 2007], RUVCorr [Freytag et al., 2015], SVA [Leek et al., 2012], and LIMMA [Ritchie et al., 2015]. For RUVCorr we used the *RUVNaiveRidge* function, which requires a set of background genes that are believed to be unrelated from treatment. For the purpose of this simulation, we selected the background genes corresponding to the nonzero entries in modules *M*_1_ … *M*_4_. For each of these methods, we corrected batch effects in gene expression, then built two co-expression matrices for case and control samples, and finally computed their difference. For COBRA, we used the case/control comparison approach described in Table 1. We also added a baseline method by subtracting the case and control co-expression networks, without any correction, and referred to this Pearson correlation difference as the “naive method”. Specifically, we computed *Ĉ*_case_ − *Ĉ*_control_, where *Ĉ*_*k*_ is the Pearson correlation computed for the samples in group *k* for *k* ∈ {case, control}. We also added a “naive batch” method that computes group differences within each batch and then averages them. Formally, we computed 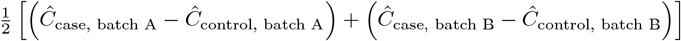 Here, *Ĉ*_case, batch A_ is the Pearson correlation computed using the case samples in batch A, and the other cases extend naturally.

We find that the distribution of co-expression values for treatment-induced gene pairs (case vs. control) and batch-induced gene pairs are confounded using the naive method without correction (Figure 2-A). However, they are more clearly separated after COBRA correction (Figure 2-B), showing that COBRA is able to better discriminate “real” treatment effects from batch effects. When comparing COBRA and naive estimates for batch and treatment effects, we see that COBRA finds a decomposition that separates each covariate of co-expression data (Figure 2-C). To quantify this improvement, we computed Receiver Operating Characteristic (ROC) curves for COBRA and the baselines we described above. In these ROC curve comparisons, we tested between real effects (positive class) against all pairs (negative class) (Figure 2-D left panel) and between real effects (positive class) against batch effects (negative class) after removing background gene pairs (Figure 2-D right panel). For the first case, COBRA does better than other methods with an AUROC of 0.96, as opposed to 0.66 for the runner-up (naive method). For the second case, COBRA’s AUROC was 0.87, while the runner-up method (naive batch) was 0.67.

**Figure 2:**
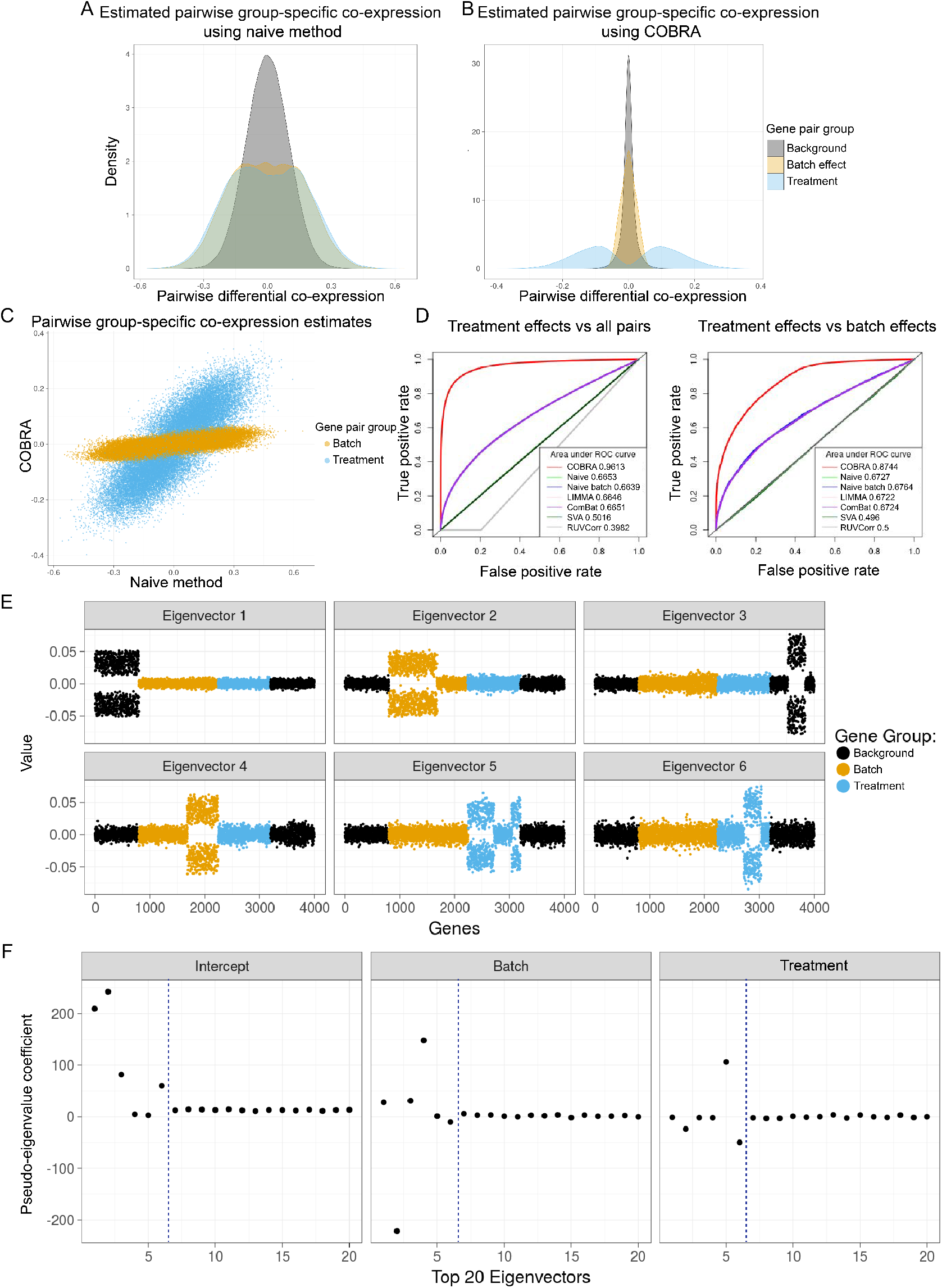
Performance of COBRA in various simulation settings. Comparison of methods for covariate-specific co-expression estimates for *in silico* data (A-C). **A. Naive differential co-expression** scores vs **B. COBRA** scores for treatment, batch, and background effects. Pearson correlation difference (naive method) failed to discriminate true co-expression compared to background, and confounded batch effects as real treatment effects. COBRA identified treatment effects in clear contrast to both background and batch effects. **C. The predicted scores of non-background gene pairs** for naive differential co-expression (x-axis) vs COBRA (y-axis) demonstrate an improved ability to separate treatment effects (blue) from batch effects (orange). **D. ROC curves** show the performance of COBRA, naive, naive batch, LIMMA, ComBat, SVA, and RUVCorr in identifying treatment gene pairs compared to all pairs (left), and in comparison to batch genes (right). Eigenvector plots show the separation of co-expression modules in simulation examples (E-F). **E. Graphical representation of the top six eigenvectors** of the original co-expression matrix plotted for all 4,000 genes. Each point is colored according to each gene’s membership in batch, treatment (case or control), or background modules. We see that eigenvectors tend to separate along with co-expression modules. **F. COBRA-computed pseudo-eigenvalues for the top 20 eigenvectors** corresponding to the three covariates (intercept, batch, treatment). Deviations from zero on the y-axis are indications of an unequal contribution of the corresponding eigenvector to the fitted co-expression estimate as shown in panel E. For example, eigenvectors 5 and 6 have nonzero pseudo-eigenvalues corresponding to the treatment variable.

We also showed that batch-induced differences in co-expression persist even after batch correction of gene expression data and complete removal of group differences (Figure S1-A). COBRA was able to remove residual batch-induced effects in co-expression for another simulation experiment where they were artificially induced (Figure S1-B).

To demonstrate that COBRA estimates are interpretable, we performed an eigendecomposition of the original co-expression matrix and plotted the top six eigenvectors for all 4,000 genes (Figure 2-E). We see that the eigenvectors tend to separate along with co-expression modules. Then, we computed COBRA’s pseudo-eigenvalues for the top 20 eigenvectors corresponding to our three binary covariates. These variables correspond to the intercept, batch, and treatment variables (case or control group). We find that COBRA assigns the largest absolute pseudo-eigenvalue coefficient (Figure 2-F) to the correct module in the original co-expression matrix (Figure 2-E). For instance, the coefficient vector corresponding to treatment has large absolute eigenvalues for components 5 and 6 (Figure 2-F), which correspond to real treatment effect gene pairs (Figure 2-E). In this case, the estimate 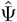 is a 3 × 4000 matrix, with the value in the *i*^*th*^ column and *j*^*th*^ row can be interpreted as the additional contribution of the *i*^*th*^ eigenvector for a one unit increase in the value of the *j*^*th*^ variable.

### 3.2 Correction of a controlled introduction of batch effects in ENCODE

We performed a benchmarking experiment to evaluate COBRA’s effectiveness in removing nonbiological differential co-expression using real-world data. We used human gene expression data [Pickrell et al., 2010], containing 153 RNA-Seq profiles for 12,424 genes in lymphoblastoid cell lines from the ENCODE project. Using the pre-processing steps described in Kuijjer and colleagues [Kuijjer et al., 2019], we considered a subset of 126 RNA-Seq profiles from 63 individuals, where each individual had been sequenced both at Yale University and at Argonne National Laboratory. Both centers used the same sequencing instrument (Illumina Genome Analyzer II), so we considered the two centers to represent two independent batches for correction.

Because there may be a batch-dependent correlation structure in the data, we constructed two groups, each containing a proportion of samples from each laboratory. We did this in fractional steps ranging from 0% of the samples being Yale to 100% of the samples being from Yale, and referred to the sample sets that contained approximately 50% of their samples from each group (either 31 or 32 since there were 63 separate research subjects) as “balanced”, and all others as “unbalanced” (Figure 3-A). We repeated each of these mixing experiments ten times and averaged the results of the measurements. For each analysis, the corresponding design matrix included three variables: intercept variable to represent average co-expression, batch variable (Yale and Argonne), and binary group variable.

**Figure 3:**
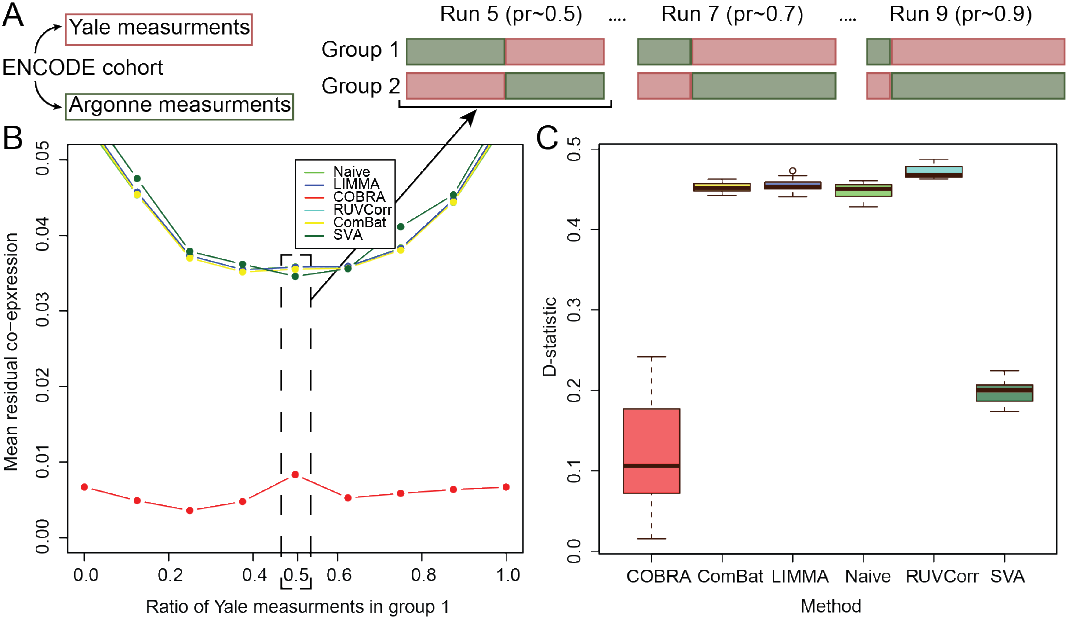
Comparison of methods for residual RNA-Seq data co-expression between centers. **A. Design of the controlled batch effect experiment.** All cohort participants were measured in both Yale and Argonne centers, and we treated each center as a batch variable. We created two groups of 63 RNA-Seq profiles in which each individual is sampled only once. We did this in fractional steps ranging from 0% of the samples being Yale to 100% of the samples being from Yale and referred to the sample sets that contained approximately 50% of their samples from each group (either 31 or 32 since there were 63 separate research subjects) as “balanced” and all others as “unbalanced”. **B. Residual co-expression between groups**. The residual co-expression (y-axis) for different proportions (x-axis) of Yale and Argonne measurements in group 1 and group 2. COBRA reduces the residuals at least by a factor of four. **C. Stability of unbalanced design (pr=0.25) in comparison to balanced design**. D-statistics for the null hypothesis of the equality of the residual co-expression’s empirical distributions for the unbalanced design (pr=0.25) in comparison to the balanced design (pr=0.5) using COBRA, ComBat, LIMMA, naive, RUVCorr, and SVA. pr: proportion.

After location-scale batch correction, we computed between-group differential co-expression analysis and because the only differences between groups were nonbiological and induced by batch, we would expect not to see group-specific co-expression. However, when we calculated the mean residual co-expression across our ten replicates at each mixing proportion, we found a significant artifactual co-expression between genes. When we applied COBRA, that residual co-expression was reduced by at least a factor of four (Figure 3-B). This finding indicates that COBRA is a more appropriate method to eliminate batch-dependent co-expression emerging in biological data, reinforcing the potential of our approach.

We also found COBRA to be more stable than the other methods across measurement proportions from each laboratory, suggesting more robust estimates. To quantify the robustness of COBRA estimates, we assessed differences in distributions between the balanced and unbalanced designs (using *pr* = 0.25 as an example unbalanced design). We selected 1000 most variable genes between lab batch “groups” and computed the Kolmogorov-Smirnov D-statistic:

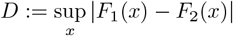

where *F*_1_ and *F*_2_ are the empirical cumulative distribution functions for the balanced and unbalanced design respectively. We estimated both cumulative distribution functions by resampling the data 1000 times to build different partitions, using both the location-scale batch corrections and COBRA. We tested for the equality of the residual group-specific co-expression’s empirical distributions using Kolmogorov-Smirnov’s test by computing ten estimates of the D-statistic (Figure 3-C), where a smaller value corresponds to weaker evidence against the null hypothesis of the distributions being equal. We observed that COBRA has smaller D-statistics than the other methods, supporting our hypothesis of being more robust. We can see that COBRA significantly outperforms the other normalization methods in removing batch-specific correlations.

### 3.3 Analysis of cancer-specific co-expression modules in thyroid cancer

We downloaded RNA-Seq human data data for thyroid cancer (THCA) from TCGA [Agrawal et al., 2014] using recount3 [Wilks et al., 2021] with sequences mapped to the gencode v26 reference gene set. For the 572 THCA samples, we considered only protein-coding genes and removed those that did not have at least one count across samples, leaving 19,711 genes. We scaled RNA-Seq counts using the *transform_counts* function from the recount3 package [Wilks et al., 2021], and normalized them using Variance Stabilizing Transformation (VST) from DESeq2 [Love et al., 2014]. Using the reported metadata we selected the following five covariates: sex (415 females, 157 males), race (381 White, 104 not reported, 54 Asian, 32 Black, 1 American Indian), stage (325 stage I, 59 stage II, 125 stage III, 2 stage IV, 51 stage IVa, 8 stage IVc, 2 not reported), batch (17 batches), age as a continuous variable (mean 47, min 15, max 89), and case/control status (the 513 cancer samples, metastatic and primary tumor, were classified as case instances (tumor); the 59 normal tissue adjacent to tumors (NAT) samples as controls). We encoded all categorical variables using the *dummy_columns* function from the fastDummies package v 1.7.3.

We computed cancer-specific co-expression networks in two ways. First, we subtracted the Pearson correlation for cancer samples and the Pearson correlation for control NAT samples. We took the absolute value or each correlation “edge” to produce a “naive network”, in which large values correspond to large differences between cancer and healthy samples. Second, we ran COBRA with a design matrix containing intercept, binary variable for cancer (0 for NAT, 1 for cancer), sex (0 for male, 1 for female), age, race, stage, and batch. All the categorical variables were coded into binary variables as described previously. After COBRA decomposition, we extracted the co-expression component corresponding to the cancer variable and took its absolute value. Since we aggregated both cancer and NAT data as input to COBRA, the decomposition decouples cancer effects from batch and other covariates, producing a cancer-specific gene co-expression network.

For both naive and COBRA co-expression networks, we used the methods in WGCNA [Langfelder and Horvath, 2008] to identify modules of correlated genes (Figure 4-A). WGCNA begins with a pairwise correlation matrix that it transforms into an adjacency matrix by raising each correlation measure to a “soft thresholding power” that is empirically determined using a scale-free topology criterion. The adjacency matrix is then transformed into a topological overlap matrix (TOM) that measures the similarity in the co-expression patterns of genes, taking into account the shared neighbors in the network. Average linkage hierarchical clustering is then performed and the resulting dendrogram is cut to identify modules; we imposed a minimum cluster size of 200 genes. We performed Gene Set Enrichment Analysis (GSEA) [Subramanian et al., 2005] for Gene Ontology (GO) [Consortium, 2019] biological process terms (Figure 4-B) and Kyoto Encyclopedia of Genes and Genomes (KEGG) [Kanehisa et al., 2017] terms (Figure S2) using *enrichGO* and *enrichKEGG* functions from the clusterProfiler package v4.10 (one-sided Fisher’s exact test, *p <* 0.05 after Benjamini-Hochberg FDR correction).

**Figure 4:**
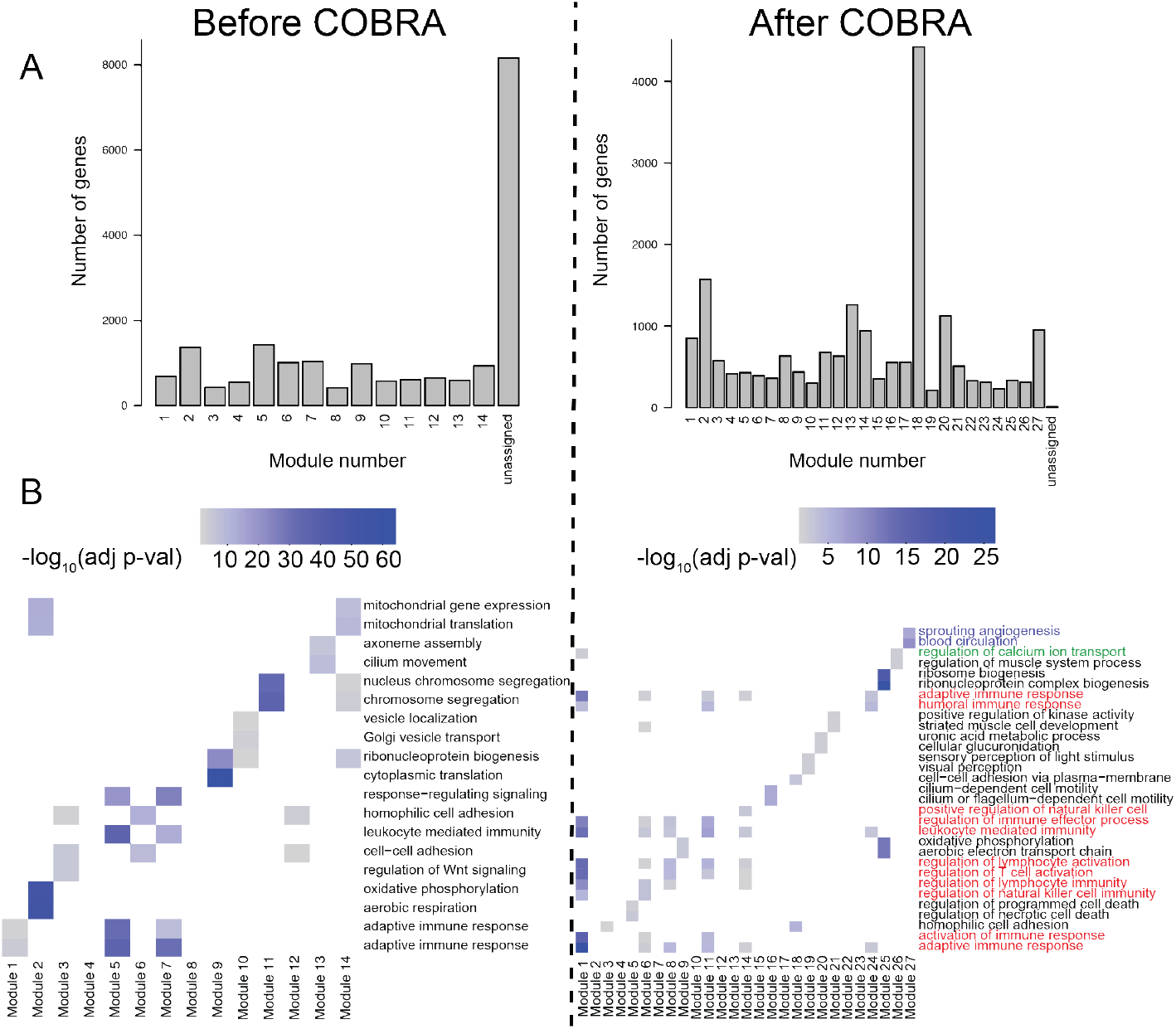
Functional modules in COBRA-corrected co-expression THCA networks. **A. Size of functional modules in co-expression networks** as computed using WGCNA before and after COBRA correction using THCA gene expression samples. Each gene is assigned to a module and some gene are unassigned. **B. Gene set enrichment analysis of functional modules** based on Gene Ontology (GO) biological process terms before and after COBRA correction. Color intensity is associated with significance levels of GO terms.

We found that COBRA networks (Figure 4-A) had a more complex community structure (27 modules) than the naive method networks (14 modules), and it had fewer unassigned genes. If we examine the cluster annotation of the two most significant terms for each module (Figure 4-B), we can find that COBRA was able to resolve the clustering of similar functional classes of genes into more precisely defined functional groups of co-expressed genes by removing residual batch effects. Of the 27 identified COBRA modules, 17 matched at least a GO term and 12 matched a KEGG term. Module 1 corresponded to cancer immune response pathways (T-cell lymphocytes and natural killer cell recruitment). Module 8 contains co-expressed genes involved in the regulation of calcium transport, which is an important physiological role of thyroid glands mediated by hormone secretion. Module 27 included genes associated with angiogenesis and blood circulation, both of which are important for tumor development and overall disease severity. KEGG modules (Figure S2) added additional evidence supporting the involvement of inflammatory processes (cytokines) for module 1, and confirmed disrupted calcium signaling in module 27. Moreover, in module 14, it further identified genes associated with osteoclast differentiation, which is induced by thyroid hormones. Enrichment analysis of the small number of modules found in the naive networks identified far less meaningful enrichment and included spurious terms such as COVID-19 and cardiomyopathy (Figure S2), quite possibly because the residual correlation structure in the non-COBRA-corrected data grouped together many unrelated genes based on spurious batch-dependent correlation.

In addition to Pearson correlation, COBRA’s design can deconvolve various association modes. Partial correlations in particular have been shown to reduce the number of spurious associations that emerge in high-dimensional co-expression networks [Shutta et al., 2023], so we used the *pcor*.*shrink* function from the corpcor package [Schafer et al., 2017] to estimate partial co-expression networks in the THCA RNA-Seq data. We then decomposed the resulting partial correlation network using COBRA, correcting for batch and the same covariates we used for Pearson correlation, and finally extracted a partial co-expression network corresponding to the cancer covariate in our design matrix. We examined the gene pairs with the 20 largest positive and negative edge scores, which included interactions between 27 genes (Figure 5-A). Among these were immune process genes such as IL16, which mediates macrophage polarization, and HTR5A, which enhances innate immunity, as well as genes such as calcitonin related polypeptide alpha (CALCA) that encodes the release of calcitonin, a hormone secreted by the thyroid gland which plays a role in calcium homeostasis.

**Figure 5:**
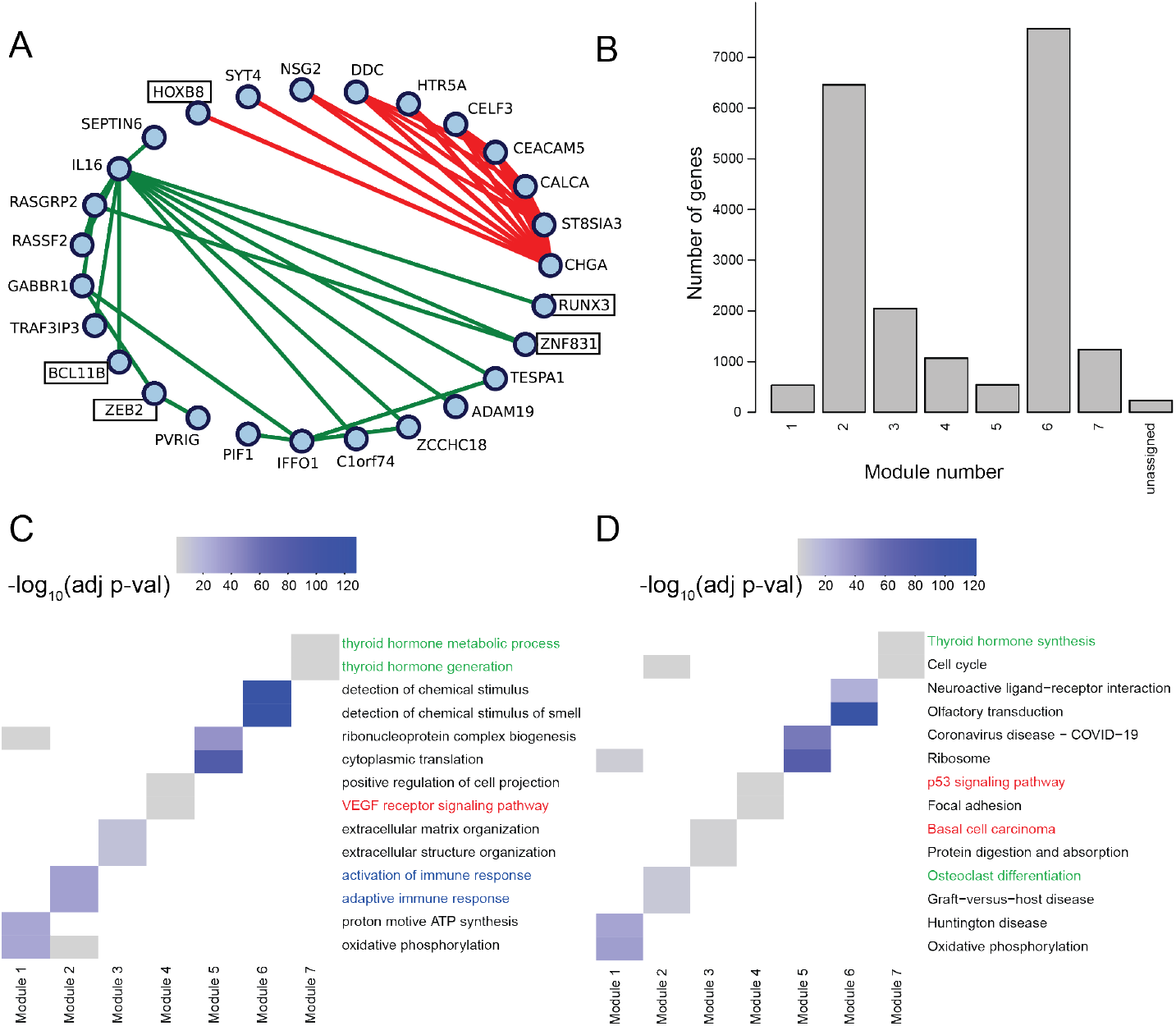
Functional modules in COBRA-corrected partial co-expression networks for THCA. **A. Largest 20 positive and negative edge weights in COBRA-corrected THCA network.** Green lines indicate positive partial correlations and red lines indicate negative partial correlations. **B. Functional modules assessment** computed using WGCNA after COBRA correction on partial correlation networks using THCA gene expression samples. **C. Gene set enrichment analysis of functional modules** in Gene Ontology (GO) biological process and **D. KEGG** after COBRA correction. Color intensity is associated with significance levels of GO and KEGG terms.

Using WGCNA, we identified seven modules in THCA partial co-expression networks, where most genes were mapped to a module (Figure 5-B), and performed GSEA selecting the two most significant terms for each module. In GO we found significant terms related to activation of immune response associated with module 2 and cancer-related signaling pathways (VEGF and p53 signaling) associated with module 4 (Figure 5-C), while in KEGG (Figure 5-D) we found terms related to thyroid physiology such as hormone synthesis in module seven and osteoclast differentiation in module 4.

### 3.4 Building accurate gene regulatory networks using COBRA batch-corrected co-expression

In our THCA co-expression analysis (Figure 5-A), we found a number of transcription factors (TFs) including BCL11B, HOXB8, RUNX3, ZEB2, ZNF831 in co-expression modules with many other genes, suggesting that some of these TFs might be regulating the expression of their correlated partners. There are a number of gene regulatory network (GRN) inference methods that use co-expression data as input [Glass et al., 2013,Khosravi et al., 2015,Van Der Wijst et al., 2018,Micheletti, 2023], and so we decided to test the effect of using COBRA batch-corrected co-expression on GRN inference. For network inference, we used PANDA [Glass et al., 2013] to infer a GRN for THCA following the same approach for other cancer types [Saha et al., 2023]. PANDA takes three prior networks as inputs: a protein-protein interaction network (PPI) indicating that some TFs can work coordinately to regulate their target genes, a prior network based on mapping TFs to their binding motifs in the genome and identifying likely TF-gene regulatory associations, and gene co-expression to capture the fact that genes deemed to be co-regulated would likely also be co-expressed. To build a PANDA THCA GRN, we first computed a COBRA covariate-specific co-expression network for cancer using cancer cases and NAT as controls, adjusting for sex, race, stage, batch, and age. We used this COBRA-corrected co-expression as an input for PANDA network inference together with PPI and motif-based prior networks downloaded from the GRAND database v1.5.4 [Ben Guebila et al., 2022a]. In the resulting GRN model, the weight of each edge quantifies the relative amount of evidence supporting the existence of a regulation relationship between the corresponding TF and gene nodes. A large positive value indicates high confidence in regulation, while a negative value indicates high confidence in the absence of regulation. Among the TFs with the greatest edge scores are members of the CEBP and POU TF families (Figure S3-A), that we suggest to be transcriptional drivers of THCA. We also conducted GSEA for genes with disrupted targeting in cancer, that is due to the increase or decrease of their regulation by TFs. To identify disrupted interactions, we computed targeting scores as the weighted in-degree for each gene, and selected high-targeted genes as those having high targeting scores, and low-targeted genes as those having low targeting scores. We find that analyzing this set of genes in OMIM database [Amberger et al., 2015] reveals similarities to genetic signatures of thyroid carcinoma (Figure S3-B,C) among other malignant diseases.

## 4 Discussion

The primary way in which gene expression has been used to study biological phenotypes has been to compare levels of gene expression between related states. This enabled to identify patterns of differential expression, which are defined by a significant difference in the mean level of expression of a gene between states. As a community, computational biologists have long recognized that effective estimation of differential expression relies on correcting for systematic differences that can arise in experimental settings, differences that are commonly referred to as “batch effects”. Consequently, it is now widely accepted that batch correction to estimate and remove those nonbiological sources of bias are essential for a robust and replicable analysis.

Despite the success of differential expression analysis, there is a growing recognition that it is not the expression of individual genes that drives phenotypic transitions, but rather changes in the expression of functional networks and pathways that define and alter cellular state. These are often found by examining patterns of correlation in gene expression data, either in a stand-alone correlation-based network analysis or as part of the process of inferring gene regulatory network models that link transcription factors to the genes they co-regulate. However, there has been no systematic study of spurious patterns of apparent co-expression arising due to batch effects, and there are no methods that specifically correct for such effects in co-expression data. COBRA addresses this gap in batch correction methods by using a matrix-factorization approach to identify and remove the effects of batch and other potentially non-interesting covariates, providing a rescaled, covariate-corrected co-expression matrix that can be used for subsequent analyses. Although COBRA was designed to model higher-order batch effects, COBRA can effectively isolate a covariate-specific co-expression.

In a simulation study, we showed that COBRA can effectively remove batch effects, including those that remain after standard batch correction. Then, using ENCODE RNA-Seq data generated on the same samples at two different sequencing centers, we were able to examine the behavior of COBRA under various batch mixing scenarios. We found that COBRA estimates were stable across different rates of imbalance, and that they differed less between balanced and unbalanced designs than did estimates from other methods. It should be noted that even for COBRA, a statistical analysis rejects the null hypothesis of both distributions being equal as p-values were greater than significance levels. This suggests there may be room for additional improvement in modeling batch-specific co-expression artifacts. In both these experiments, we benchmarked COBRA against other batch correction methods, and it is important to highlight the conceptual differences in our approach. For all other methods, we corrected for batch effects in gene expression before computing co-expression networks, while COBRA corrects for batch effects after computing co-expression, which led to a more effective correction of higher-order batch effects.

In our analysis, we also found that while linear combinations of covariates-associated co-expression exactly reconstruct the original co-expression matrix, there is no guarantee that the domain of individual components will be realistic. Specifically, some COBRA co-expression entries may lie outside the [−1, 1] interval used to define the range of possible correlation values. This concern is somewhat mitigated, however, in the analysis of thyroid cancer (THCA) that we performed as the COBRA co-expression scores had, at most, 5.9% entries falling outside the expected range (Table S1).

In analyzing data from THCA, we constructed correlation networks, partial correlation networks, and gene regulatory networks comparing tumor and normal samples. We showed that COBRA facilitates the discovery of meaningful biological patterns of co-expression, and that it can be used as part of a GRN inference workflow. For example, in the THCA COBRA partial co-expression network (Figure 5-A), we took the 20 highest-scoring positively and negatively correlated edges. Among those we found seven TF-gene associations involving five TFs. Given that TFs represent only a small fraction (< 10%) of protein-coding genes, this is far more than we would expect by chance.

Using COBRA-corrected co-expression data from the same THCA study, we demonstrated that COBRA estimated batch-corrected, covariate-specific co-expression matrices that can be effectively used to estimate and compare GRNs between conditions. As both COBRA and PANDA are available through netZoo [Ben Guebila et al., 2023], we modified PANDA to allow COBRA correction to be incorporated into the analysis when the user supplies the parameter *cobra_design_matrix*. We found that the CEBP (CCAAT/enhancer-binding protein) and POU (Pit-Oct-Unc) families of transcription factors play important roles in THCA. In particular, the POU domain transcription factor family includes proteins that are key to developmental processes such as embryonic pluripotency and neuronal specification. In general, the POU transcription factors are not typically associated with thyroid cancer. OCT1/POU2F1, the only widely expressed POU factor, operates on target genes associated with proliferation, immune modulation, and more recently identified targets like oxidative and cytotoxic stress resistance and metabolic regulation. Oct1 is pro-oncogenic in multiple contexts and has prognostic and therapeutic value in various epithelial tumor settings, including thyroid cancer [Vázquez-Arreguín and Tantin, 2016]. Further, the TF SOX12, which shows increased expression in thyroid cancer tissue and cells, and promotes cell proliferation, migration, invasion, and tumor growth, is reported to interact with the POU family. SOX12 knockdown inhibits the expression of POU2F1, POU3F1, and POU2F1. POU3F1 can reverse the effects of SOX12 knockdown on thyroid cancer cells, indicating their role in the progression of thyroid cancer [Su et al., 2022].

We also modified additional netZoo methods to integrate with COBRA, such as DRAGON, a method to infer multimodal partial correlation networks, by specifying the *method* argument in COBRA to *dragon*, in addition to *pearson* for Pearson correlation and *pcorsh* for shrunken partial correlation networks. We also plan to provide COBRA as a pre-processing correction step in other netZoo methods [Chen and Padi, 2022, Micheletti, 2023].

COBRA represents a significant step forward in batch correction methodology, allowing more effective removal of batch-dependent correlation structures while at the same time providing a way of identifying covariate-specific patterns of co-expression to be identified. The availability of the code and various tutorials as online Jupyter notebooks in [Ben Guebila et al., 2022b] will facilitate the use of this method.

## Code and data availability

COBRA is available in R and Python through the netZoo packages [Ben Guebila et al., 2023] netZooR v1.5 and netZooPy v0.9.16 (https://netzoo.github.io). We also provide tutorials in R and Python through Netbooks (http://netbooks.networkmedicine.org/) [Ben Guebila et al., 2022b]. The code and data for the analysis are available at the GitHub repository (https://github.com/QuackenbushLab/cobra-experiments). Human RNA-Seq data from the ENCODE project was downloaded from GEO under accession number GSE19480.

