## Supplementary information document for "Higher-order correction of persistent batch effects in correlation networks"

### Supplementary material

#### Contents


**1 Optimization** **1**

**2 Supplementary methods** **2**

2.1 RNA-seq data analysis with controlled batch effects . . . . . . . . . . . . . . 2

2.2 Benchmark setting . . . . . . . . . . . . . . . . . . . . . . . . . . . . . . . . . 4

**References** **6**


### 1 Optimization

COBRA optimizes the following objective function (Equation 1 in the main text):

$$\arg\min_{\boldsymbol{\Psi}} ||\boldsymbol{C} - \frac{1}{n}\sum_{i=1}^{n} \boldsymbol{Q} diag(\boldsymbol{X}_i^T \boldsymbol{\Psi}) \boldsymbol{Q}^T||_F^2, \tag{1}$$

where $\boldsymbol{C}$ is the sample covariance matrix of a zero-centered gene expression $\boldsymbol{G}$, and $\boldsymbol{Q}$ is the matrix of the eigenvectors corresponding to non-zero eigenvalues in the eigendecomposition $\boldsymbol{C} = \boldsymbol{Q}\boldsymbol{D}\boldsymbol{Q}^T$. The goal is estimating the matrix $\boldsymbol{\Psi} \in \mathbb{R}^{k \times r}$ ($r$ being the number of non-zero eigenvalues of $\boldsymbol{C}$), quantifying the impact of the $q$ covariates in the design matrix on each eigenvalue. We now show that a global optimum for Equation 1 can be computed in closed-form. The first crucial observation is that, since $\boldsymbol{C}$ is symmetric, then $\boldsymbol{Q}$ is orthogonal. We get

$$\arg\min_{\boldsymbol{\Psi}} ||\boldsymbol{C} - \frac{1}{n}\sum_{i=1}^{n} \boldsymbol{Q} diag(\boldsymbol{X}_i^T \boldsymbol{\Psi})\boldsymbol{Q}^T||_F^2$$

$$\arg\min_{\boldsymbol{\Psi}} ||\frac{1}{n}\sum_{i=1}^{n} \boldsymbol{G}_{\cdot,i}\boldsymbol{G}_{\cdot,i}^T - \frac{1}{n}\sum_{i=1}^{n} \boldsymbol{Q} diag(\boldsymbol{X}_i^T \boldsymbol{\Psi})\boldsymbol{Q}^T||_F^2 \quad \text{Definition of sample covariance}$$

$$\arg\min_{\boldsymbol{\Psi}} ||\frac{1}{n}\sum_{i=1}^{n} \left(\boldsymbol{G}_{\cdot,i}\boldsymbol{G}_{\cdot,i}^T - \boldsymbol{Q} diag(\boldsymbol{X}_i^T \boldsymbol{\Psi})\boldsymbol{Q}^T\right) ||_F^2 \quad \text{Commutativity}$$

$$\arg\min_{\boldsymbol{\Psi}} ||\frac{1}{n}\sum_{i=1}^{n} \left(\boldsymbol{Q}^T\boldsymbol{G}_{\cdot,i}\boldsymbol{G}_{\cdot,i}^T\boldsymbol{Q} - diag(\boldsymbol{X}_i^T \boldsymbol{\Psi})\right) ||_F^2 \quad \text{Orthogonal matrices preserve the Frobenius norm}$$

$$\arg\min_{\boldsymbol{\Psi}} \sum_{h=1}^{r} \left(\frac{1}{n}\sum_{i=1}^{n} \left(\boldsymbol{Q}_{\cdot,h}^T\boldsymbol{G}_{\cdot,i}\boldsymbol{G}_{\cdot,i}^T\boldsymbol{Q}_{\cdot,h} - \boldsymbol{X}_i^T \boldsymbol{\Psi}_{\cdot,h}\right)\right)^2 \quad \text{Optimizing } \Psi \text{ only affects diagonal elements}$$

Since the terms above are positive and each one corresponds to a different column of $\hat{\boldsymbol{\Psi}}$, we can decouple the optimization and solving for each $h \in [r]$ separately:

$$\hat{\boldsymbol{\Psi}}_{\cdot,h} = \arg\min_{\boldsymbol{\Psi}} \left(\sum_{i=1}^{n} \boldsymbol{Q}_{\cdot,h}^T\boldsymbol{G}_{\cdot,i}\boldsymbol{G}_{\cdot,i}^T\boldsymbol{Q}_{\cdot,h} - \boldsymbol{X}_i^T \boldsymbol{\Psi}_{\cdot,h}\right)^2 \tag{2}$$

In other words, we are minimizing the sum of the residuals for a standard linear regression problem. Here, every unbiased estimator achieves the optimum in Equation 1, and we pick the maximum-likelihood estimator (MLE) because it is the candidate solution with lowest variance. Assuming that the design matrix has an intercept column, the MLE has closed-form solution:

$$\hat{\boldsymbol{\Psi}}_{\cdot,h} = (\boldsymbol{X}^T\boldsymbol{X})^{-1}\boldsymbol{X}^T \left[\boldsymbol{Q}_{\cdot,h}^T\boldsymbol{G}_{\cdot,i}\boldsymbol{G}_{\cdot,i}^T\boldsymbol{Q}_{\cdot,h}\right]_{i=1}^{n}. \tag{3}$$

### 2 Supplementary methods

#### 2.1 RNA-seq data analysis with controlled batch effects

We used gene expression data from ENCODE [5], containing 153 RNA-seq profiles for 12,424 genes in lymphoblastoid cell lines. Using the pre-processed data described in Kuijjer and colleagues [3], we considered a subset of 126 RNA-seq profiles from 63 individuals, where each individual has been sequenced both at Yale University and at Argonne National Laboratory. Both centers used the same technology (Illumina Genome Analyzer II) which should reduce batch effect due to sequencing technology. For our purpose, the two centers represent the two batches to consider for correction. We constructed 2 groups each containing a proportion of samples from each centre. We considered the proportion 0.5 to model a balanced design by taking 31 (49%) measurements from Yale and 32 (51%) measurements from Argonne, forming a partition of about 0.5 each. Since each group contains RNA-seq

Table S1: Statistics of COBRA components

| **Dataset**/ Component | Ratio of entries not in $[-1,1]$ | Minimum entry | Maximum entry |
|---|---|---|---|
| **Thyroid carcinoma** | | | |
| $\bar{C}$ | 0.90% | -1.51 | 1.60 |
| Cancer | 0% | -0.87 | 0.87 |
| Batch 1 | 0% | -0.40 | 0.55 |
| Batch 2 | 0% | -0.42 | 0.59 |
| Batch 3 | 0.15% | -1.07 | 0.74 |
| Batch 4 | 0% | -0.48 | 0.76 |
| Batch 5 | 0% | -0.35 | 0.82 |
| Batch 6 | 0% | -0.71 | 0.77 |
| Batch 7 | 0% | -0.53 | 0.53 |
| Batch 8 | 0% | -0.22 | 0.43 |
| Batch 9 | 0% | -0.38 | 0.67 |
| Batch 10 | 0% | -0.41 | 0.29 |
| Batch 11 | 0% | -0.29 | 0.47 |
| Batch 12 | 0.45% | -1.13 | 1.02 |
| Batch 13 | 0% | -0.24 | 0.19 |
| Batch 14 | 0% | -0.37 | 0.36 |
| Bath 15 | 0% | -0.37 | 0.47 |
| Batch 16 | 0.32% | -1.12 | 0.73 |
| Batch 17 | 0% | -0.57 | 0.61 |
| **ENCODE (0.5 vs 0.25 proportion)** | | | |
| $\bar{C}$ | 0.0008% | -0.92 | 1.16 |
| Group | 0% | -0.19 | 0.16 |
| Batch | 0% | -0.32 | 0.25 |

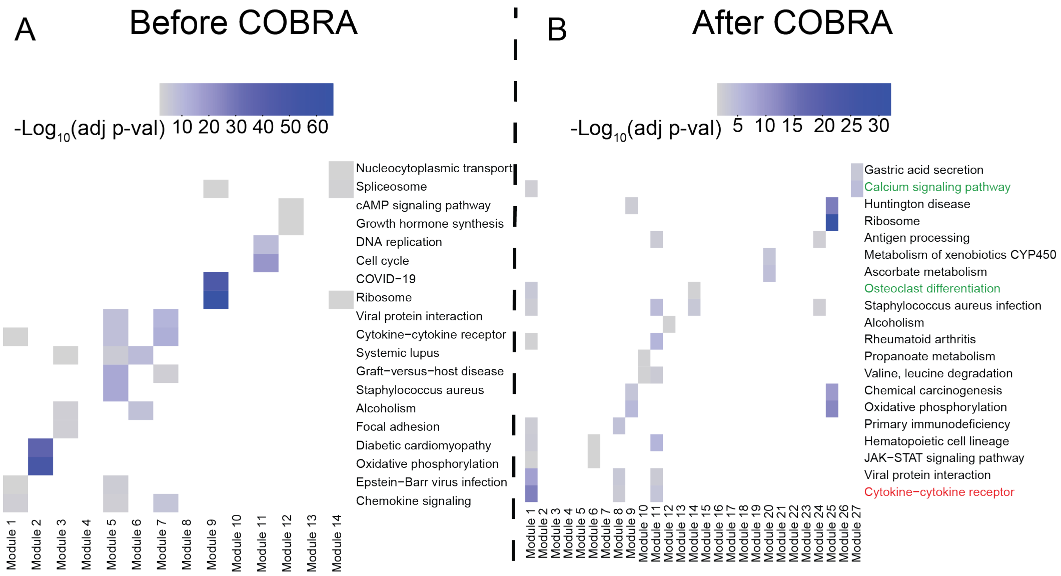


Figure S1: **Gene set enrichment analysis in KEGG database of thyroid cancer networks.** Analyses were done before COBRA correction **(A)** and after COBRA correction of co-expression networks **(B)**. Color intensity is associated with significance levels of KEGG term enrichment.

assays on the same 63 individuals, one would expect that, following batch correction, there would be minimal differential expression and co-expression between groups. Our design matrix for COBRA included three variables: intercept to represent average co-expression, batch (Yale and Argonne), and case (group1 and group2). We extracted the co-expression matrix corresponding to the case variable as our batch-corrected COBRA co-expression matrix. To assess differences in distributions between groups, we selected 1000 most variable genes and computed the Kolmogorov-Smirnov D-statistic:

$$D := \sup_x |F_1(x) - F_2(x)|$$

where $F_1$ and $F_2$ are the empirical cumulative distribution functions for the balanced and an unbalanced design with 0.25 measurement proportion. We estimated both cumulative distribution functions using 1000 samples.

#### 2.2 Benchmark setting

In simulated and real-world data settings, we benchmarked COBRA against the following batch correction methods: ComBat [2], RUVCorr [1], SVA [4], and LIMMA [6]. For RUVCorr we used the *RUVNaiveRidge* function, which requires a set of negative control genes. In our simulation study we picked the background genes, while in the real-world settings we picked the 60 genes corresponding to the 30 entries with smallest absolute co-expression. We also assessed differential co-expression by subtracting the case and control co-expression networks to build a differential network and refer to this Pearson difference as ”naive method.”

Formally, we computed $C_{\text{case}} - C_{\text{control}}$, where $C_k$ is the Pearson correlation computed for the samples in group $k$ for $k \in \{\text{case, control}\}$. For simulated data, we additionally added a "naive batch" method which computes group differences within each batch and then averages them. Formally, we computed $\frac{1}{2}\left[(C_{\text{case, batch A}} - C_{\text{control, batch A}}) + (C_{\text{case, batch B}} - C_{\text{control, batch B}})\right]$. Here $C_{\text{case, batch A}}$ is the Pearson correlation computed using the case samples in batch A, and the other cases extend naturally. Note that in our simulation settings there are exactly two batches.
